# Feeding disruptions lead to a significant increase in disease modules in adult mice

**DOI:** 10.1101/2023.11.07.565937

**Authors:** Xiaoqin Mou, Pengxing Nie, Renrui Chen, Yang Cheng, Guang-Zhong Wang

**Author notes:** Corresponding author; (G.W.).

## Abstract

Feeding disruption is intimately associated with numerous diseases, yet its molecular mechanisms remain to be elucidated in circadian biology. We discovered that in the liver of adult mice, feeding disruption can lead to a substantial increase in disease modules in the transcriptome. This increase in disease modules occurs far earlier than the individual’s aging and disease onset. In contrast, calorie restriction significantly reduces the formation of disease modules. This offers a critical missing link between feeding disruption and the molecular systems mechanisms of disease.

## Main

Circadian rhythm disruption can affect the expression of a multitude of genes, leading to negative health consequences ^1-3^. Such disruption can exacerbate the severity of diseases, such as increasing the risk of neurological, psychological, and immune-related diseases ^1,3^. Among these, feeding disruption is associated with the risk of numerous diseases, such as cardiovascular and metabolic disorders ^4^. Conversely, calorie restriction and time-restricted eating, can offer several benefits, including delaying aging and extending lifespan ^5-8^. However, the systemic molecular mechanisms underlying these effects are yet to be fully elucidated.

Disease genes often interact with each other, forming a module of pathogenic genes ^9,10^. The emergence of these modules heightens the organism’s susceptibility to the related disease ^9-11^. We hypothesis that under feeding disruption conditions, disease genes are likely to exhibit synchronized expression patterns, and form more disease modules. To assess this hypothesis, we examined dietary data from various disruption scenarios. Additionally, we employed various disease datasets to validate these observations and delve into the potential molecular system-level mechanisms associated with feeding disruption.

We compared three groups of feeding disruptions in healthy adult mice: ad libitum feeding (AL) versus calorie restriction with evenly distributed food intake (CR-spread), calorie restriction with feeding restricted to 2 hours during the day (CR-day-2h) versus 2 hours during the night (CR-night-2h), and calorie restriction with feeding restricted to 12 hours during the day (CR-day-12h) versus 12 hours during the night (CR-night-12h) ^12^. Our initial analysis focused on a set of 22 diseases (DS22) identified in mice under these conditions (Table S1) ^12^.

To determine the significance of these disease modules, we conducted 1,000 random permutation tests, repeating each experiment 20 times ^9^. Under the CR-spread condition, we noted the significant formation of modules associated with six diseases in the mouse liver’s circadian transcriptome (Figure 1A and table S2), including Lymphoma (p <= 0.002) and Fibrosis (p <= 0.004). However, under the AL condition, we observed the formation of disease modules related to an additional six diseases (Figure 1B and table S2), such as Hemangiosarcoma (p <= 0.007), Hemangioma (p <= 0.001), and Adenoma (p <= 0.008). This was a significantly higher number compared to the disease modules formed under the CR-spread condition (Figure 1C, p = 4.15×10^−9^, Wilcoxon rank-sum test). These findings suggest that calorie restriction alone, even without time restriction, can significantly reduce the number of disease modules in the liver.

**Figure 1.**
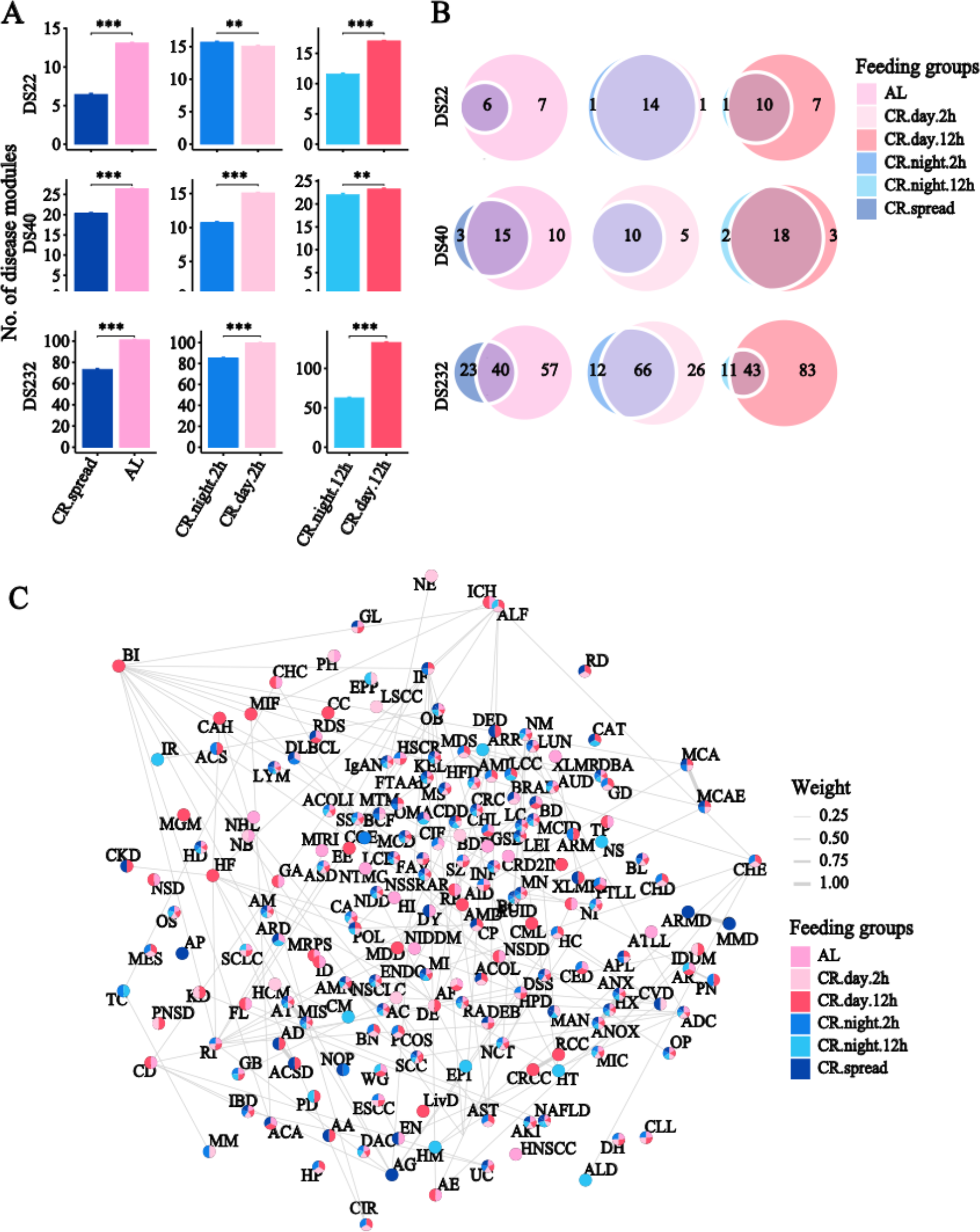
Significant increase in disease modules under feeding disruption in adult mice. A. Comparison of the number of disease modules detected under different feeding conditions. Each bar represents a specific feeding condition and the height of the bar indicates the number of disease modules under that condition. The bar charts represent the number of disease modules for three disease datasets (DS22, DS40, DS232) across six feeding conditions: AL, CR-spread, CR-day-12h, CR-night-12h, CR-day-2h, and CR-night-2h. The significance levels are denoted as follows: *** for p < 0.0001, ** for p < 0.001 in the Wilcoxon rank-sum test. B. Venn diagrams illustrating the overlap of disease modules for DS22, DS40, and DS232 across different feeding experiments. C. Network representation of the enriched disease modules in the six feeding experiments. The node labels refer to specific diseases (abbreviated). Nodes are color-coded based on the feeding conditions (AL, CR-spread, CR-day-2h, CR-night-2h, CR-day-12h, and CR-night-12h). Lines connecting nodes represent significant associations between the diseases or conditions, with the thickness of the line indicating the weight, as defined by the Jaccard coefficient between two disease modules.

Similarly, under the CR-night-12h condition, we detected 11 disease modules (Figure 1X), such as Lymphoma (p <= 0.006), Fibrosis (p <= 0.007), Hemangioma (p <= 0.042), Adenoma (p <= 0.019), and Thrombus (p <= 0.0036). Conversely, under the CR-day-12h condition, there was 17 disease modules detected (Figure 1A), including Hemangiosarcoma (p <= 0.001), Cyst (p <= 0.004), Hyperplasia (p <= 0.014), and Histiocytic sarcoma (p <= 0.009). This resulted in a significantly higher number of disease modules compared to those formed under the CR-night-12h condition (Figure 1C, p = 2.86×10^−9^, Wilcoxon rank-sum test, see Tables S2).

To validate the findings between feeding disruption and disease modules, we employed two additional disease sets. The first set included diseases previously reported to be related to circadian disruption (DS40) ^4^. The second set was compiled from the DisGeNET database, encompassing gene sets for all disease types and resulting in a collection of 232 diseases (DS232) (Table S1) ^13^. We replicated the permutation experiments and discovered that, in the DS40 disease dataset, the mice under AL condition had 7 more significant disease modules compared to the CR-spread condition (p = 1.51×10^−8^, Figure 1A). Likewise, the mice under CR-day-2h condition had 5 additional disease modules compared to the CR-night-2h condition (p = 3.73×10^−9^, Figure 1B). The CR-day-12h condition had one more significant disease module than the CR-night-12h condition (p = 0.00039). In the DS232 disease dataset, the AL condition had 34 more significant disease modules compared to the CR-spread condition (average disease modules: 97 vs. 63, p = 2.96×10^−8^), the CR-day-2h condition had 14 additional potential disease modules compared to the CR-night-2h condition (average disease modules: 92 vs. 78, p = 3.046×10^−8^), and the CR-day-12h condition had 72 additional potential disease modules compared to the CR-night-12h condition (average disease modules: 126 vs. 54, p = 3.072×10^−8^) (Figure 1B and Table S2). Despite the mice appearing healthy under all these conditions, their potential risk for diseases markedly increased under disrupted conditions. This significant rise in disease motifs suggests that feeding disruption exerts a substantial influence on the health of adult mice.

Does feeding disruption under conditions without calorie restriction also lead to an increase in disease motifs? Our comparison of the liver circadian transcriptome between unrestricted feeding (ALF) and time-restricted feeding (TRF) conditions ^14^ in healthy adult mice showed that the average number of detected disease motifs across the three disease module datasets (DS22, DS40, and DS232, Figure 2A) was 8, 12, and 76 in the TRF conditions, respectively. In contrast, under the ALF condition, the average number of detected disease modules were 10, 17, and 87, displaying a significant increase in each disease dataset (Wilcoxon test p = 3.716×10^−6^; p = 2.968×10^−8^; p = 1.539×10^−9^) (Figure 2A, Tables S3). These findings further validate the observations made under calorie-restricted conditions.

**Figure 2.**
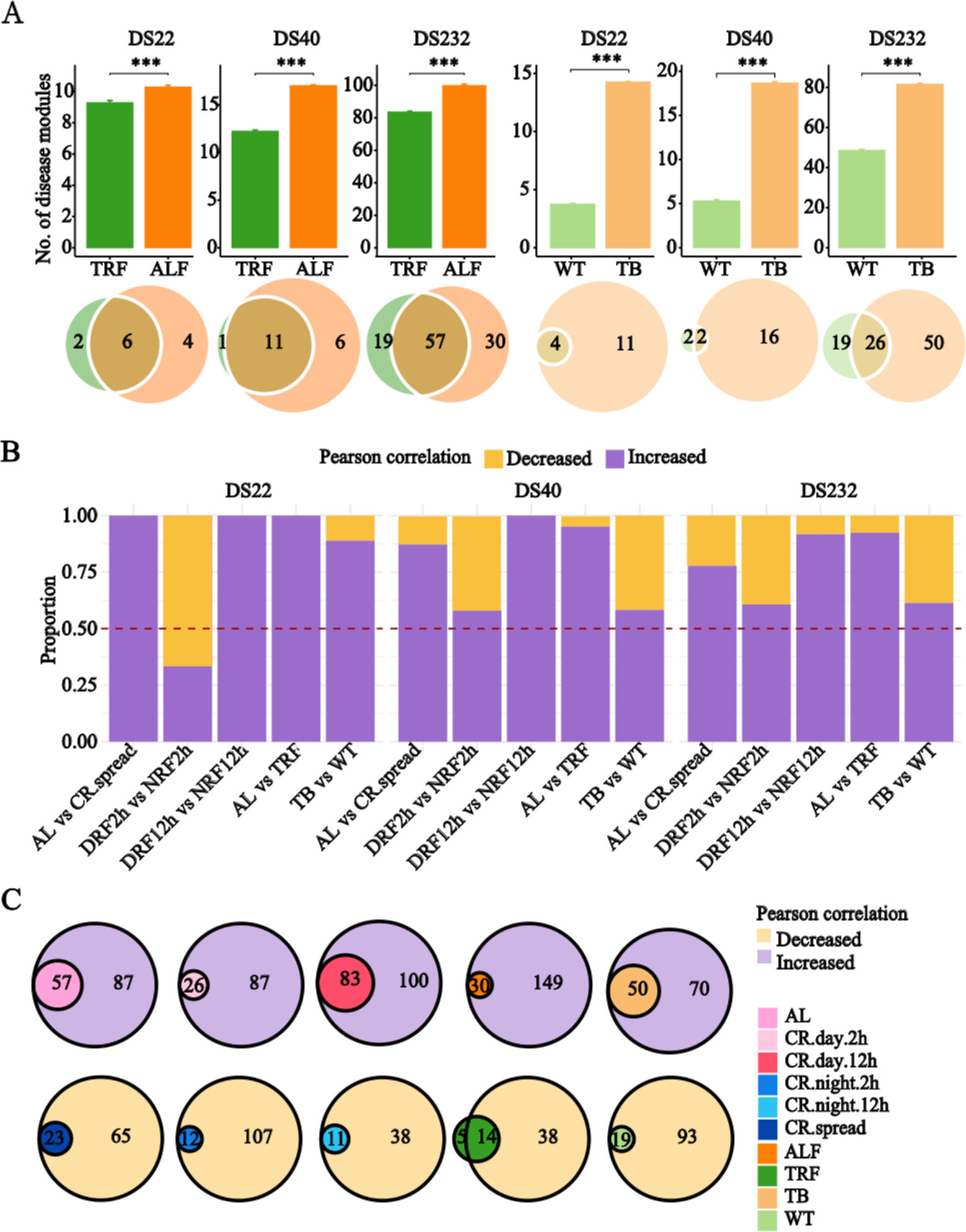
Relationship between the increase in the number of disease modules and their elevated co-expression levels. A. Bar charts illustrating the number of disease modules across different conditions: TRF compared to ALF, and WT compared to TB. Accompanying Venn diagrams below the bar charts depict the overlap of detected disease modules among the experiments. Significance levels are denoted by asterisks: *** indicates p < 0.0001 in the Wilcoxon rank-sum test. B. Proportion of disease modules with significantly increased co-expression under disrupted conditions compared to controls. Experimental conditions, including comparisons such as AL vs TRF and AL vs CR-spread, are displayed on the x-axis. The color purple indicates disease modules with increased correlation under disrupted conditions, while gold signifies disease modules with decreased correlation under these conditions. The red dashed line serves as a reference, indicating a proportion of 0.50. CR-day-12h, DRF12h; CR-night-12h, NRF12h; CR-day-2h, DRF2h; and CR-night-2h, NRF2h. C. Significant disease modules typically exhibit high co-expression. Overlapping circles represent the interplay between the number of significant disease modules and those with high Pearson correlation coefficients under each condition. Each condition is distinctly color-coded, as detailed in the legend located at the bottom right.

Further, studies on lung cancer have demonstrated that these diseases can impact the activity of the liver transcriptome by disrupting pathways such as AKT, AMPK, and SREBP ^15^. In our study, we observed that in mice with lung adenocarcinoma, the number of disease modules in liver tissue rose by 175%, 350%, and 68.9% across the three sets of disease modules respectively (Figure 2A and Tables S4). This further indicates that the number of disease modules in the liver significantly increases under disease conditions.

We next investigated the functions of the disease modules formed under disruption conditions. For this, we constructed WGCNA co-expression analysis networks and performed enrichment analysis on the disease module genes ^16^ Disease genes under AL conditions were significantly enriched in WGCNA modules turquoise (P = 0.00011), and lightyellow (P = 2.87×10^−6^). Disease genes under CR-day-12h conditions were significantly enriched in WGCNA modules magenta (P =1.09×10^−18^), salmon (P = 0.0005),and lightyellow (P =0.0383) (Figure S1 and Table S5). These modules were significantly enriched in inflammation and immune-related biological pathways, such as activation of immune response (P = 8.34×10^−29^), endolysosomal toll-like receptor signaling pathway (P = 0.000249), and inflammasome complex assembly (P = 0.00201) in AL (Figure S2, Table S6). Conversely, disease genes under non-disrupted conditions were enriched in WGCNA modules related to biological pathways such as metabolic process and other basic biological pathways (Figure S2, Table S6). These findings align with the results of Acosta-Rodríguez et al ^12^.

Is the increase in the number of disease modules due to the increased co-expression of these disease genes? To answer this question, we compared the co-expression magnitude between disease genes under feeding disruption and controls. Although an increase in co-expression of disease genes does not necessarily lead to the formation of significant disease motifs, we found that the co-expression of genes in most disease modules significantly increased under disrupted conditions, as shown in Figure 2B and 2C. Moreover, under the AL condition, the number of disease modules in the liver exhibiting increased co-expression was more than three times the number of those showing decreased co-expression (1 vs. 21, 6 vs. 32, 52 vs. 179, Figure 2B and Table S7). This indicates that the formation of disease modules is intricately linked to the elevated co-expression of disease genes.

Our results suggest that feeding disruption has a significant impact on the transcriptome of healthy adult individuals. Genes affected by dietary disruption undergo substantial changes in expression, which can strongly influence the transcriptome and potentially make them disease-related genes. Simultaneously, at the module level, dietary disruption substantially increases the number of potential disease modules within the circadian transcriptome. This rise in the number of disease modules during prolonged disruption significantly elevates the likelihood of individuals developing diseases. Importantly, the formation of these disease modules occurs significantly earlier than the onset of these diseases in individuals. These discoveries provide a framework for understanding the systematic mechanisms of disease development in circadian biology.

## Methods

### Data Sources

We utilized multiple liver circadian transcriptome datasets in this study. The data on calorie-restricted feeding experiments in healthy mice were obtained by subjecting 6-week-old male C57BL/6J mice to various methods of calorie restriction (CR) ^12^. This included ad libitum feeding (AL), providing 70% of baseline food intake within two hours at the beginning of the day (CR-day-2h) and night (CR-night-2h), feeding the mice continuously every 90 minutes during the day for 12 hours (CR-day-12h) and night (CR-night-2h), as well as feeding the mice continuously every 160 minutes (CR-spread). At 6 months of age, the mice were released into a continuous dark environment while maintaining the feeding schedule. Sample collection began 36 hours after the onset of continuous darkness, and liver samples were collected every 4 hours for RNA sequencing experiments. Two replicates were collected each time over a 48-hour period, with 24 mice under each feeding condition, totaling 144 samples (4). The data for non-calorie-restricted feeding in healthy mice came from a study that fed 12-week-old C57BL/6J mice under either AL or TRF conditions for 7 weeks ^14^. Liver tissue samples from two mice were taken every 2 hours over a 24-hour period for RNA-seq sequencing. The dataset for the lung tumor bearing model (TB) in mice^15^ included microarray data from liver samples collected from wild-type mice and lung tumor metastasis model mice at ZT 0, 4, 8, 12, 16, and 20. Each time point had three replicates, resulting in a total of 36 samples.

### Data processing

For the RNA-seq data obtained from the mouse calorie-restricted feeding experiments, which comprised 21,869 genes and 288 samples. We selected 13,997 genes where the count number was at least 3 in a minimum of 24 samples. Similarly, For the RNA-seq data from the AL and TRF dataset under normal conditions, which includes 40,677 genes across 96 samples. We selected 15,115 genes that were expressed (count number > 0) in at least half of all samples. This led to an approximately equal number of genes in both datasets. All RNA-seq data samples then underwent a variance-stabilizing transformation (VST) using the ‘vst’ function in the DESeq2 package ^17^ --. The microarray data from the lung tumor bearing model in mice was normalized using the ‘rma’ function in the oligo package (version = 1.64.1) ^18^.

### Identification of disease modules

Three disease sets were assembled for this study. The first disease set encompassed 22 diseases closely linked to tumors and inflammation ^12^, while the second included 40 diseases that have previously been reported to be associated with circadian disorders ^4^. The third disease set consisted of 232 diseases obtained from the DisGeNET (v7.0) database(website: https://www.disgenet.org/)^13^. In each dataset, we filtered genes that exhibited expression in at least half of the samples for subsequent identification of significant disease modules.

To identify significant disease modules ^9,19^, we first computed the average Pearson correlation coefficient for all the disease genes associated with each specific disease. we then randomly selected an equivalent number of genes from all genes, calculating the Pearson correlation coefficient among these randomly selected genes. This permutation experiment was repeated 1000 times. We then compared the average correlation coefficient of the disease genes with that of the randomly selected genes to calculate a p-value. If the p-value was less than 0.05, we considered that we had detected a significant disease module. Each disease module calculation was repeated 20 times in total. Finally, we compared the differences in the number of significant disease modules under different experimental conditions (case vs. control).

### Weighted gene co-expression network analysis (WGCNA)

We conducted weighted co-expression gene module analysis using the “blockwiseModules” function from the WGCNA package (13). We set the soft threshold power to 16, the maximum module size to 13,168, used Pearson’s method for correlation calculation, set the network type to signed, the TOM type to signed, the minimum module size to 30, and the threshold for merging modules to 0.25. These settings enabled the identification of unsupervised WGCNA modules formed under the specified experimental conditions. Subsequently, we conducted a one-tailed Fisher’s test on the disease genes to determine whether they are significantly associated with specific WGCNA modules.

### Functional enrichment analysis

In this study, we utilized the clusterProfiler package ^20^(Version = 4.8.2) within the R environment to conduct Gene Ontology (http://geneontology.org/) enrichment analysis, specifically focusing on Biological Processes (BP). Subsequently, we applied the Benjamini-Hochberg correction method for multiple testing adjustments. In order to further visualize the results of the enrichment analysis, we employed the ‘dotplot’ function to generate bubble plots in R.

### Statistical Analysis

All co-expression coefficients were calculated using the Pearson correlation method, as computed by the ‘cor’ function in R. Wilcoxon rank-sum tests were used for the detection of differences between groups and Fisher’s test was used for enrichment analysis. A p-value < 0.05 was considered statistically significant. All analyses were performed using R version 4.3.1, and visualizations were created using ggplot2 v.3.4.2.

## Code availability

The primary codes used for the analysis can be found in the supplementary code materials.

## Aacknowledgements

This work is supported by grants from the National Natural Science Foundation of China (Grant Nos. 81827901 and 32170567) to G-Z, W.

## Ddeclaration of interests

The authors declare no competing interests

